# *In vitro* liquid-liquid phase separation induced by respiratory syncytial virus proteins and RNA

**DOI:** 10.1101/2025.11.25.690368

**Authors:** Vincent Basse, Tanushree Agarwal, Tomas Sneideris, Charles-Adrien Richard, Joris Troussier, Jean-Jacques Vasseur, Françoise Debart, Jean-François Eléouët, Ella de Csillery, Tuomas Knowles, Marie Galloux

## Abstract

Respiratory syncytial virus (RSV) causes severe respiratory infections, with viral replication occurring in cytoplasmic membraneless viral factories formed by liquid-liquid phase separation (LLPS). The interactions between the RSV nucleoprotein N, phosphoprotein P, transcription factor M2-1, and RNA drive these condensates. Here we employed a microfluidic Phase Scan platform, together with biochemical and cellular assays to systematically characterise LLPS involving RSV proteins and RNA. We identified optimal stoichiometric ratios of N oligomers and P tetramers for condensate formation, demonstrated that monomeric N inhibits LLPS, and revealed that M2-1 enhances condensate formation by increasing multivalency. Notably, we discovered that M2-1 preferentially binds 5’ capped RNA, distinguishing it from N, which binds uncapped RNA. These findings elucidate molecular determinants of RSV viral factory assembly and subcompartmentalisation, providing insights into viral replication mechanisms and informing potential antiviral strategies targeting LLPS processes.

## Introduction

Liquid-liquid phase separation (LLPS) is a key process in the formation of a wide variety of cellular organelles that are involved in diverse cytoplasmic and nuclear pathways and mechanisms (*1*). Viruses, which are obligate intracellular pathogens, were also shown to induce the formation of functional organelles in host cells, relying on LLPS. More specifically, the transcription and replication of viruses belonging to the *Mononegavirales* (*MNV*) order, which possess a single-strand negative-sense RNA genome (*2*), take place inside cytoplasmic membraneless viral factories (VF) that display liquid-like properties (*3*). These viro-induced organelles facilitate the concentration of viral proteins and nucleic acids, as well as host factors required for the viral polymerase activity or involved in the host innate immune response (*4*–*6*).

Within *MNV*, the respiratory syncytial virus (RSV) is the main cause of bronchiolitis in young children worldwide (*7*), resulting in over 100,000 infant deaths every year (*8*). RSV is also responsible for lower respiratory tract infections (LRTI) in the elderly and immunocompromised individuals (*9*). In the past two years, three vaccines have been approved for the elderly and pregnant women (*10*–*13*). Although vaccinating pregnant women indirectly protects newborns during their first weeks of life, the only effective treatment currently available for children is a preventive injection of monoclonal antibodies that target the fusion protein F, which is responsible for virus entry into host cells (*14*, *15*). Despite these important advances in effective prophylactic treatments, they are only expected to provide short-term protection. In the absence of effective therapeutics, a better characterisation of the viral cycle and the identification of novel targets thus remain necessary. The formation of VF, which relies on highly specific viral protein-protein interactions (PPIs), is a key step in the viral cycle that could be targeted for antivirals development.

The RSV genome, of 15.2 kb, is composed of 10 genes encoding 11 proteins, with the M2 gene displaying two ORFs that encode two proteins (M2-1 and M2-2 proteins) (*16*). The viral genome is constantly enwrapped by the nucleoprotein (N), forming left-handed non-canonical helical nucleocapsids (NC) (*17*, *18*). Shortly after viral entry by the fusion of viral and cellular membranes, the ribonucleoprotein complex (RNP), composed of the NC in association with the polymerase L, its cofactor the phosphoprotein P, and the transcription factor M2-1, is released into the cytoplasm where VF are formed. These organelles can be detected by microscopy 8 hours post-infection, and were shown to be dynamic, their size increasing throughout the infection and events of fusion and fission being observed (*4*). This dynamic depends on transient, low-affinity PPIs essential for L polymerase function, particularly the P-mediated recognition of NCs. The P protein is also involved in the recruitment of the viral M2-1 protein and the cellular protein phosphatase 1 (PP1) into VF (*19*, *20*). Interestingly, a switch in the phosphorylation state of M2-1, mediated by PP1, was shown to induce its accumulation with newly synthesised viral mRNA into liquid sub-compartments of VF, called IBAGs (for inclusion bodies associated granules) (*4*, *20*). These observations suggest that the functioning of RSV VF depends on a fine-tuned temporal and spatial regulation of their structural organisation. Mechanistically, co-expression of only N and P proteins in eukaryotic cells was shown to allow the formation of pseudo-VF (*21*). *In vitro* reconstitution assays further revealed that N-RNA oligomers and P tetramers are the main drivers of pseudo-VF formation through LLPS (*22*). The P protein, considered as a hub in RNPs, is composed of 241 residues and presents three structural domains: a central domain of tetramerization (P_OD_, 131 to 151), and two intrinsically disordered regions (IDRs), i.e., the N-terminal (P_NTD_) and C-terminal domains (P_CTD_) (**Figure 1a**). Through diverse PPIs, P_NTD_ mediates the recruitment of M2-1 and PP1 to VF (*19*, *20*), but also acts as a molecular chaperone that stabilises the neosynthesised N in its monomeric and RNA-free form (N^0^), competent for subsequent specific encapsidation of the viral antigenomes and genomes (*23*, *24*). On the other hand, P_CTD_ interacts with L through multiple contacts, and with the NC (*23*, *25*–*27*). Consistent with their respective role in the binding of N-RNA oligomers and P tetramerization, P_CTD_ and P_OD_ were shown to be required for pseudo-VF morphogenesis (*22*). The N protein, of 391 residues, folds into N- and C-terminal globular domains (N_NTD_ and N_CTD_ respectively) separated by a hinge region involved in RNA binding, and presents flexible N- and C-terminal arms that play a critical role in N oligomerization (**Figure 1a**) (*28*). Although only NCs serve as template for the L protein, different N-RNA oligomers co-exist in infected cells, among which 10-or 11-N oligomers complexed to RNA that form N rings, also observed in virus filaments (*18*, *28*, *29*). Finally, the M2-1 protein is composed of 194 amino acids and forms tetramers. This protein displays a N-terminal zinc-binding domain (ZBD) identified as an RNA binding site, a tetramerization domain, a core domain that binds both RNA and P in a competitive manner, and a C-terminal tail (*19*, *30*–*32*). Although the binding domains of P with N and M2-1 and the role of RNA in N and M2-1 activity are well characterised, their role in VF formation and organisation remain poorly understood.

**Figure 1.**
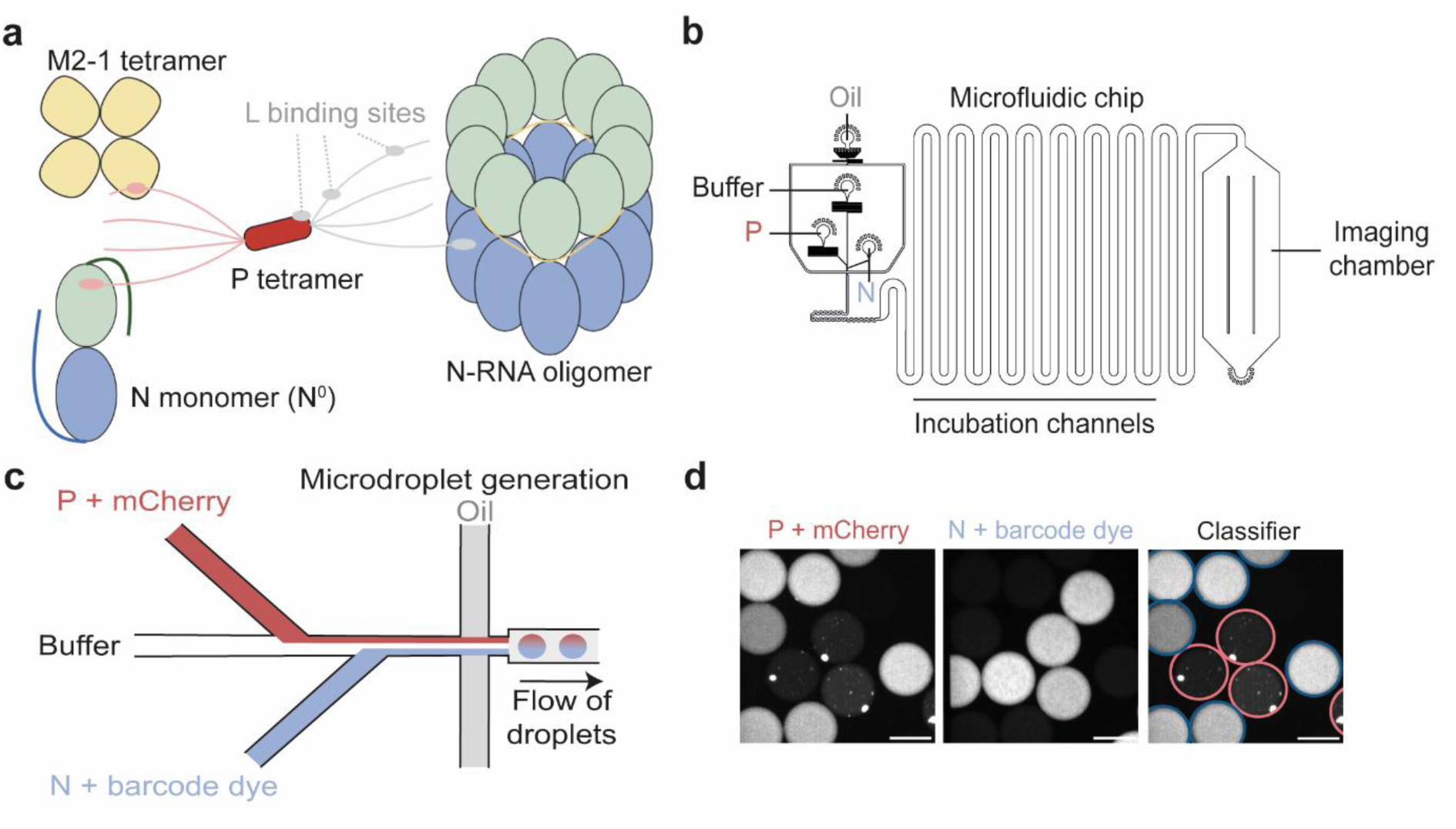
RSV proteins of interest and the Phase scan method. **(a)** The interactions of P tetramer with N-RNA oligomers (represented as N-rings), the monomeric and RNA-free N^0^, and the M2-1 tetramer are represented. The binding sites of L are indicated. N and P subdomains are highlighted with different colours: green for N_NTD_, blue for N_CTD_, pink for P_NTD_, red for P_OD_, grey for P_CTD_. The yellow thin line within the N-RNA oligomer represents encapsidated genomic/antigenomic RNA. Made with *Biorender.com*. **(b)** Microdroplets with varying concentrations of N and P proteins are generated using a microfluidic device controlled by automated pressure pumps. **(c)** Prior to droplet formation, aqueous protein solutions are combined under laminar flow conditions at the droplet generation junction. The resulting microdroplets are then imaged using fluorescence microscopy. **(d)** Fluorescence microscopy images show P protein tagged with mCherry and N protein with Alexa488 within the microdroplets. Droplets are classified as either phase-separated (outlined in red) or homogeneous (outlined in blue), based on the presence or absence of visible condensates. Scale bars represent 100 μm.

The goal of this study was to gain information on the role of the RSV N, P and M2-1 proteins, as well as RNA, in LLPS events associated with the formation of RSV condensates *in vitro*. Using the Phase Scan platform, we monitored LLPS events with various combinations of RSV proteins and nucleic acids. Our results provide a detailed characterisation of the respective roles of P and N in pseudo-VF formation and also highlight the impact of M2-1 on LLPS. We further reveal a specific interaction between M2-1 and 5’ capped RNA, shedding light on a new element that could contribute to the specificity of interaction of M2-1 with mRNA and N with the viral genome. Altogether, our findings reveal key elements that could further explain the structural organisation and dynamics of RSV VF.

## Results

### Design of Phase scan experiments

To construct high-density phase diagrams of RSV protein condensates and explore a broad chemical space, we here employed the Phase Scan platform (*33*) (**Figure 1**), using a constant buffer (20 mM Tris-HCl pH 7.4, 150 mM NaCl buffer) without any crowding agent. All the recombinant proteins used in this study were produced in *E. coli* and purified using methods already published (**Figure S1**) (*19*, *22*, *23*, *34*). In our system, the purified recombinant wild type (WT) N protein corresponds to N-RNA rings (*28*), and the N-P40 construct, which consists in the fusion of N to the 40 N-terminal residues of P (P40), was used to mimic the monomeric RNA-free N^0^ protein (*34*, *35*). To directly observe LLPS through the formation of fluorescent condensates, each experiment was performed with one of the components fused to a fluorescent probe, i.e., either N or P proteins fused to the mCherry, or Cy5 labelled RNA. Noteworthy, to minimise the potential impact of mCherry on LLPS, considered labelled proteins were always mixed to an excess of WT proteins (1:10). Microdroplets were generated by systematically modulating the flow rates of aqueous inputs – N, P, M2-1, RNA, and buffer, while maintaining a constant flow of HFE-7500 fluorinated oil containing 1.2% (w/w) polyglycerol-based triblock surfactant (**Figure 1b**). All flows were precisely regulated by automated pressure-based microfluidic pumps. Within the microfluidic chip, the laminar flow ensured the aqueous streams remained separated until encapsulation, allowing for the formation of droplets with well-defined and varied compositions. Fluorescence microscopy was used to image the droplets in flow (**Figure 1b-d**). Each protein component was tagged with a unique fluorescent barcode, enabling precise quantification of its concentration within each droplet. Barcode intensities were compared to calibration curves generated from known barcode concentrations, enabling detailed mapping of phase behaviour across a wide range of conditions. Droplets were categorised as either phase-separated or mixed based on the presence or absence of visible condensates. The resulting raw image data were compiled into phase diagrams, where component concentrations were plotted against phase separation status. Scatter plots were generated to visualise the estimated probability of phase separation across the concentration space, providing a comprehensive landscape of condensate behaviour.

### Minimal requirements for LLPS involving the RSV N and P proteins

Previous work established that RSV pseudo-VF morphogenesis is driven by the interactions between N-RNA oligomers and P tetramers, both in cells and *in vitro* (*22*). These heterotypic condensation events take place in a stoichiometrically tuned manner, where optimal P_tetramer_:N_rings_ ratios for liquid droplet formation ranged from 5:1 to 5:8, and down to sub-micromolar concentrations (*36*). While LLPS was reported to be observable without crowding agents, most experiments were conducted in the presence of Ficoll or PEG 4000 (*22*, *36*–*38*). Here, we characterised the heterotypic condensation of N and P in the absence of any crowding agents. At first, a combination of phase scans, either using a mix of mCherry-P/WT-P (ratio 1:10) together with N rings, or the other way around, were performed. Phase diagrams showed that LLPS occurs within a specific window of stoichiometric P:N ratios. Analysed in terms of monomers, LLPS requires a minimum ratio of 1 P per 6 N, and ceases when P becomes excessively abundant (exceeding 5:1). Normalisation of these monomer ratios to the oligomeric states of both proteins (P tetramers and N decamers) indicates that optimal phase separation occurs at P_tetramer_:N_rings_ ratios ranging from a minimal threshold of approximately 0.5:1 to 1:1 and up to 3:1 to 12:1 (**Figure 2a**). Crucially, the boundaries of this LLPS zone are highly dependent on the absolute N concentration, defining a much narrower window at higher protein loads. The lower limit represents the minimal concentration of P required for nucleation and assembly, while the upper limit demonstrates a re-entrant phase behaviour where excess P inhibits condensation. Importantly, we confirmed that the condensates were spherical and that neither mCherry-P/WT-P nor mCherry-N-/WT-N formed condensates alone, serving as a critical negative control (**Figure S2a, b**).

**Figure 2.**
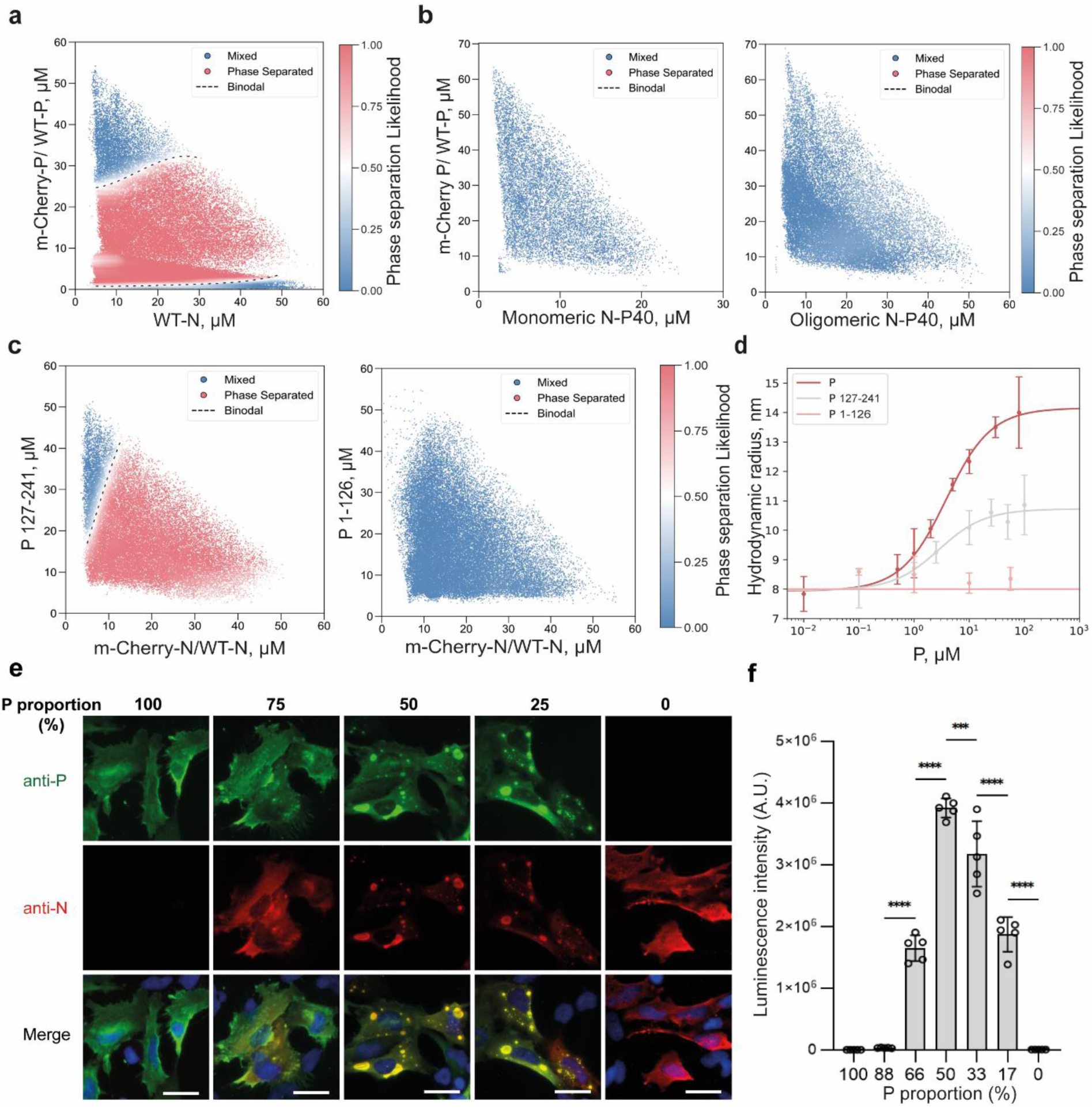
Characterisation of N/P LLPS. **(a)** Phase diagram depicting mCherry-P/WT-P versus WT-N protein concentrations. Red points correspond to phase-separated droplets, while blue points indicate homogeneous ones. The dashed line depicts the phase boundary (acting as visual reference). **(b)** Analysis of the capacity of mCherry-P/WT-P and monomeric N-P40 (left) or oligomeric RNA-free N-P40 (right) protein to induce LLPS. **(c)** Study of the propensity of P_127-241_ (left) or P_1-126_ (right) to phase separate with mCherry-N/WT-N. The scatter plots display the approximate likelihood of phase separation across a spectrum of concentrations of RSV N and P proteins. The number of distinct microdroplets examined were **(a)** n = 158874, **(b)** n = 14802 (left) n = 44734 (right), **(c)** n = 53368 (left), n = 56442 (right). **(d)** Hydrodynamic radius (R_H_) of N at 0.5 µM with varying P, P_127-241_, and P_1-126_ concentrations. Errors represent standard deviations from triplicate measurements. At high concentrations of P and P_127-241_, a plateau is reached, indicating saturated binding with N. No binding was observed with P_1-126_. **(e)** Varying quantities of plasmid DNA encoding N and P were co-transfected in BEAS2B cells. Cells were fixed 24 post-transfection and labelled with anti-P (AF488, green) and anti-N (AF594) antibodies, and pseudo-VFs were observed by fluorescence microscopy. Nuclei were stained with Hoechst 33342. Bars, 10 μm. **(f)** BSRT7/5 cells were transfected with plasmids encoding viral proteins M2-1 and L, varying quantities of plasmid DNA encoding N and P, a plasmid encoding the pMT/Luc minigenome as well as the pCMV-βGal for transfection standardisation. Cells were lysed 24h post-transfection and viral RNA synthesis was quantified by measuring the luciferase activity. Each luciferase minigenome activity value was normalised based on β-galactosidase expression. The graph corresponds to one representative experiment out of three independent experiments (**Figure S3**) performed in 5-plicate. Error bars represent standard deviation (±SD) calculated based on one representative experiment. Data were analysed using a one-way ANOVA followed by multiple comparisons test. Asterisks indicate the level of statistical significance: (***) p < 0.001; (****) p < 0.0001; “ns”, not significant.

To further investigate the molecular requirements for LLPS, we also tested the contributions of RNA and specific domains of P. In agreement with previous findings, no phase separation was observed when using monomeric, RNA-free N-P40 with P (**Figure 2b**) (*34*). Interestingly, this was also true at high N-P40 concentrations known to induce N-P40 oligomerisation (**Figure 2b**) (*34*), emphasising the critical importance of RNA for LLPS. We then confirmed that the N-terminal domain of P (P_1-126_) did not induce LLPS with N rings (**Figure 2c**), whereas the protein deleted from this domain (P_127-241_) did induce phase separation (**Figure 2c**) (*22*). Interestingly, an excess of P_127-241_ was less limiting for LLPS than full-length P, suggesting that deletion of the N-terminal part of P increases its propensity for phase separation. We therefore assessed N-P binding affinities using micro diffusional sizing (MDS) by measuring the hydrodynamic radius (R_H_) of N rings as the concentration of P was increased. The R_H_ increased from a plateau to a larger radius, indicating complex formation and yielding a dissociation constant (K_d_) of 3.3±0.3 µM (**Figure 2d**). As expected, no increase in R_H_ was reported when P_1-126_ was added to N rings, indicating that no binding occurred. In contrast, increasing concentrations of P_127-241_ led to an increase in R_H_, confirming its binding to N rings (K_d_ = 2.6±0.9 µM). While no significant difference in N binding affinity between full-length P and P_127-241_ was observed, these micromolar K_d_ values are consistent with the transient, low-affinity PPIs known to favour LLPS and with previously published data (*23*).

In parallel, we assessed the importance of the P/N ratio for pseudo-VF formation in cells. Cells were transfected with different ratios of plasmids encoding P and N, and the presence of pseudo-VF was monitored by fluorescence microscopy after immunolabelling. We observed that while a strong excess of P (≥ 75%, corresponding to a P_plasmid_:N_plasmid_ ratio of ≥ 3:1) impaired cytoplasmic condensates formation, suggesting a P-induced re-entry of phase, an equivalent proportion of both proteins or an excess of N (up to 75% compared to P, i.e., a 1:3 ratio) allowed phase separation (**Figure 2e**). We then evaluated the relationship between LLPS and polymerase activity in cells, using a functional minigenome assay (*25*). Our results showed a direct correlation between the optimal P/N ratio for LLPS and those for efficient viral polymerase activity (**Figure 2f**). Due to this functional assay’s reliance on the co-transfection of 6 plasmids in cells, some variability in polymerase activity was reported between experiments (**Figure S3**). However, excess of P constantly had a negative impact on the polymerase activity, in contrast to N overexpression. This cellular inhibition by P excess aligns remarkably with the re-entrant phase behaviour observed in our *in vitro* phase diagrams, which showed that high P_monomer_:N_monomer_ ratios lead to the inhibition of LLPS. The cellular data confirms that the functional window of the viral polymerase complex is tightly restricted by the abundance of P. A disparity in P:N ratios was, however, observed between *in vitro* and *in cellula* experiments. Such differences are likely due to the heterogeneity of N oligomers in cells where NC can form, and/or to the impact of cellular co-factors interacting with P and N, which dynamically modulate the LLPS equilibrium. Overall, our observations confirm the importance of the P/N ratio for LLPS in both systems. Our results also confirm the capacity of N-RNA rings and P tetramers to induce LLPS in the absence of crowding agents and further suggest that the highly disordered N-terminal domain of P acts as a modulator, altering the propensity for phase separation *in vitro*.

### Monomeric RNA-free N-P40 alters LLPS formation

As previously mentioned, the newly synthesised monomeric N^0^ protein must be recruited to VF during infection for new viral genomes and antigenomes encapsidation. Supporting this, we have recently shown that monomeric N-P40-GFP is recruited to pseudo-VF formed in cells in the presence of WT N and P proteins (*35*). However, the precise impact of this recruitment on LLPS remains to be characterised. We therefore set out to monitor the effect of adding monomeric N-P40 on *in vitro* LLPS events observed upon co-incubation of N rings and mCherry-P/WT-P. We observed that the gradual addition of N-P40 led to a striking reduction in LLPS between N rings and P (**Figure 3a**). Further increase of N-P40 correlated with a dose-dependent decrease in mCherry condensates, with no more N/P condensates forming above 10 µM of N-P40, even with higher concentrations of N rings and/or P (**Figure 3a**). While we cannot entirely rule out that fusion of the P40 peptide to monomeric N might have introduced steric hindrance that impaired N^0^/P interactions, these observations suggest that monomeric N-P40 competes with N rings for P binding *in vitro*, thereby inhibiting N/P LLPS. It is noteworthy that while P is known to bind N-RNA oligomers and N^0^ through P_CTD_ and P_NTD_ respectively, it remains unclear whether one P tetramer can bind both forms of N simultaneously. We then investigated further N-P interactions using band shift analysis of proteins migration on agarose native gels (*34*). As shown on figure 3b, P, N rings, and N-P40 each display a specific migration profile. Co-incubation of N rings and P led to the observation of a single band specific to the N-P complex (**Figure 3b**). In contrast, upon co-incubation of P and N-P40, the bands corresponding to each unbound protein were observed, and only a minor additional band corresponding to a P-N-P40 complex was detected. These data highlight the poor capacity of P to interact with N-P40 and correlate with phase scan data, showing the absence of LLPS in our conditions.

**Figure 3.**
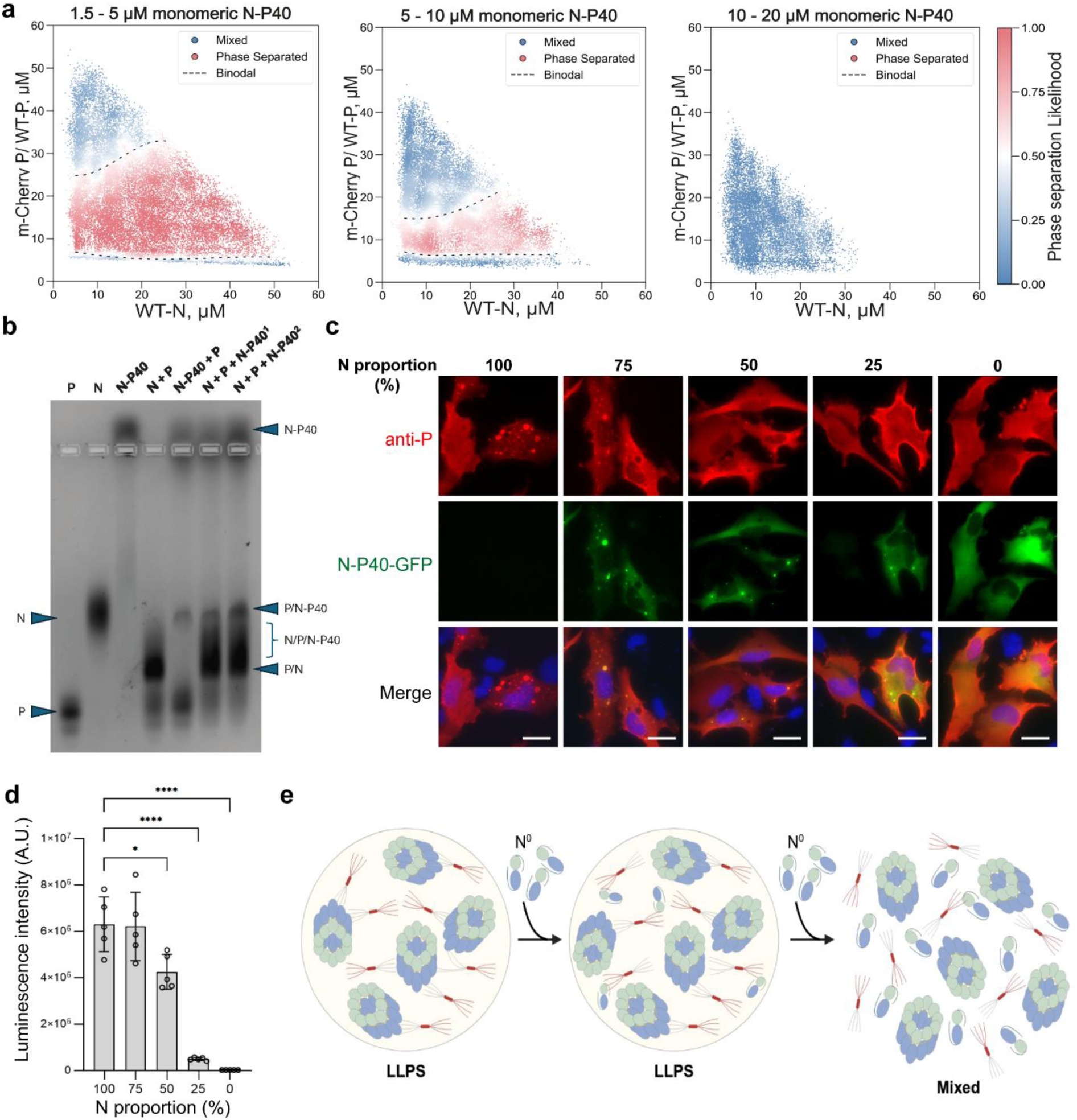
Impact of monomeric N-P40 on N/P LLPS and viral polymerase activity. **(a)** Phase Diagrams obtained in the presence of mCherry-P/WT-P and N rings with increasing concentrations of N-P40. The scatter plot displays the approximate likelihood of phase separation. Red and blue points indicate phase separated and mixed regions respectively. The dashed line depicts the phase boundary. The number of distinct microdroplets examined was n= 40612 (left), n= 45445 (centre), n = 36551 (right). **(b)** Analysis of N, P, N-P40 and N-P complexes by agarose native gel. Bands corresponding to the individual proteins or complexes are indicated. **(c)** A plasmid encoding P and varying quantities of plasmid DNA encoding N and N-P40-GFP were co-transfected in BEAS2B cells. Cells were fixed 24 post-transfection and labelled with anti-P (AF594, red) antibody, and pseudo-VFs were observed by fluorescence microscopy. Nuclei were stained with Hoechst 33342. Bars, 10 μm. **(d)** BSRT7/5 cells were transfected with plasmids encoding P, M2-1, L, pMT/Luc minigenome, pCMV-βGal, and varying quantities of N and N-P40. Cells were lysed 24h post-transfection and viral RNA synthesis was quantified by measuring the luciferase activity. Each luciferase minigenome activity value was normalised based on β-galactosidase expression. The graph corresponds to one representative experiment out of three independent experiments (**Figure S3**) performed in 5-plicate. Error bars represent standard deviation (±SD) calculated based on one representative experiment. Data were analysed using a one-way ANOVA followed by multiple comparisons test. Asterisks indicate the level of statistical significance: (**) p < 0.01; (****) p < 0.0001. **(e)** Schematic representation of how the recruitment of N^0^ to N/P VF must be kept under a certain threshold, otherwise LLPS – and thus polymerase activity – is impaired. Made with *Biorender.com*.

When then co-incubating N rings, P, and monomeric N-P40, the band corresponding to P-N-P40 complex was still observed, together with a smeary lower band that could correspond to the N-P complex mixed to different N-P-N-P40 complexes. Interestingly, when increasing the proportion of N-P40, the intensity of the unbound N-P40 and P-N-P40 complex bands increased without altering the N-P-N-P40 smear intensity. This observation correlates with our phase scan data, providing biochemical clues that the binding of monomeric N-P40 to P might compete with the LLPS of N rings and P.

In parallel, we also assessed the impact of N-P40 overexpression in cells on the capacity to induce LLPS. Cells were transfected with a constant quantity of the plasmid encoding for P, together with various ratios of N/N-P40-GFP encoding plasmids. Here, only P was immunolabelled before observation by fluorescence microscopy of P and N-P40-GFP localisation. As previously described, co-expression of equivalent amounts of N and N-P40 (or lower amount of N-P40) with P allowed us to observe pseudo-VF similar to those obtained in the presence of N and P, where N-P40 was recruited (**Figure 3c**). However, an excess of N-P40 led to a clear defect in the formation of cytoplasmic inclusions. Of note, no small inclusions where P and N-P40 could co-localise were observed compared to previous published data, obtained in a different cell line (*35*). We then performed minigenome assays to evaluate the impact of monomeric N-P40 on the viral polymerase activity. Cells were transfected with different proportions of plasmids encoding N or N-P40, together with the plasmids required for minigenome expression. While the polymerase activity was not affected in the presence of a low amount of N-P40, up to 25% of the total N protein pool (**Figure 3d, S3**), a sharp decrease in activity occurred with higher N-P40 proportions, leading to a total loss of activity for the highest N-P40 quantities. Such alterations directly mirror the capacity to form pseudo-VF in cells.

Altogether, our results show that while monomeric N-P40 binds P with a low affinity, its excess strongly impairs N/P LLPS *in vitro* (**Figure 3e**). These findings underscore the importance of a tight regulation of the N^0^ pool during infection, as its concentration appears to be a critical factor in the formation of VF (*34*, *35*).

### M2-1 participates in pseudo-VF morphogenesis

During infection, M2-1 is recruited to VF by P (*4*, *19*). We first investigated the capacity of P and M2-1 to induce LLPS in the absence of N (**Figure 4a**). Phase scans conducted with mCherry-P/WT-P and M2-1 confirmed their propensity to form condensates (**Figure 4b**) (*37*). Our results also showed that while small concentrations of P (10 μM) limited LLPS, an overabundance of M2-1 tetramers compared to P tetramers did not disrupt condensate formation (**Figure 4b**). Importantly, M2-1 alone did not form condensates, confirming that the phase separation observed was driven by heterotypic interactions (**Figure S2c**). These results suggest that although the recruitment of M2-1 to VF in cells depends on its interaction with the P_NTD_, LLPS between P and M2-1 likely involves additional, transient contacts between other domains of both proteins (*37*, *39*). We next explored how M2-1 impacts the LLPS of the core N/P components. *In vitro* co-incubation of N, P and M2-1 was previously shown to induce the formation of three-component condensates (*37*, *38*). We found here that the gradual addition of M2-1 to N/P profoundly enhanced the overall propensity for LLPS (**Figure 4c**). This effect was similar at first in magnitude to the enhancement observed upon deletion of P_NTD_ (**Figure 2c**), and then even stronger as M2-1 concentration increased. However, we cannot rule out that these condensates correspond to both P/M2-1 and N/P/M2-1 LLPS.

**Figure 4.**
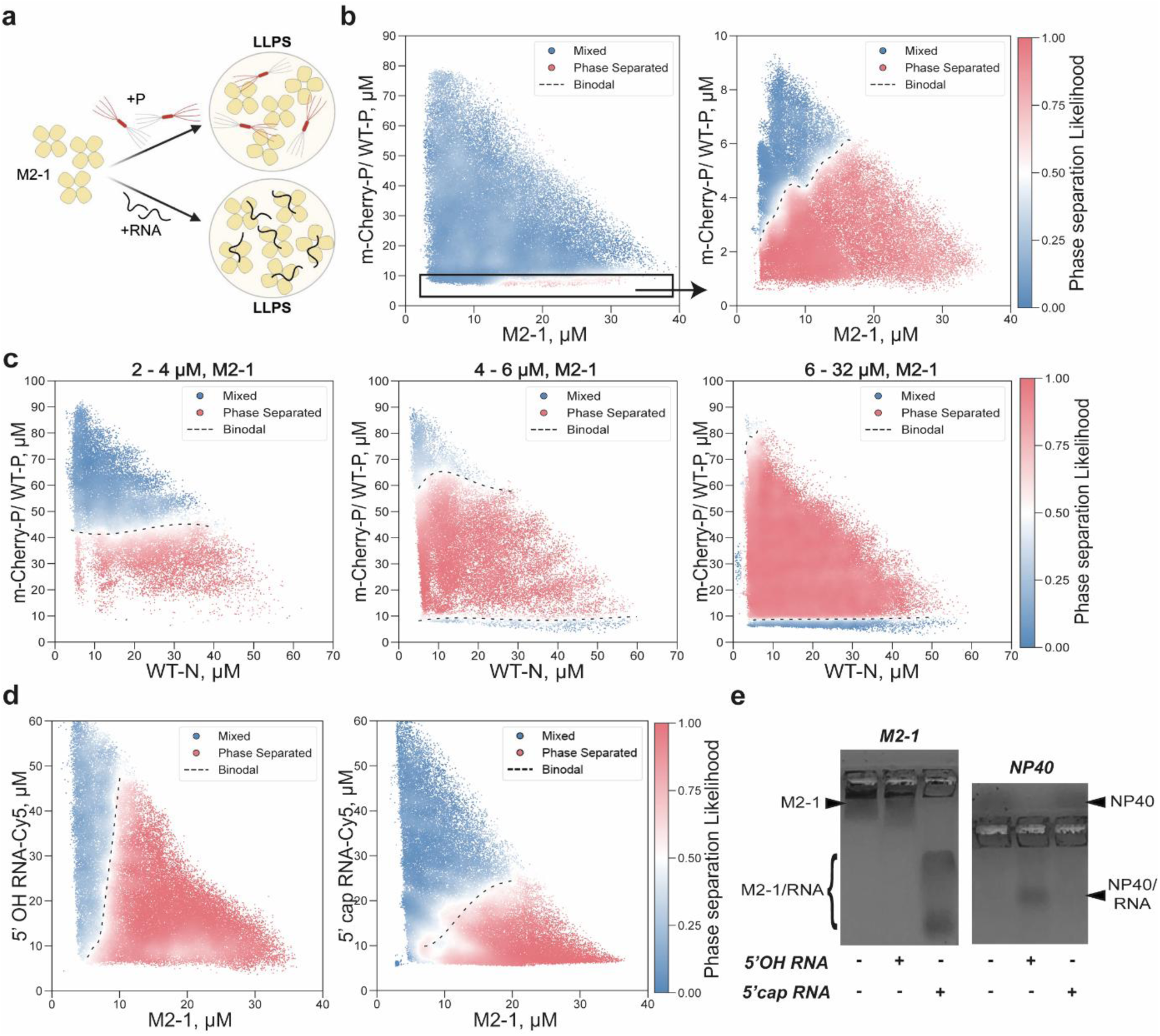
M2-1 actively participates in pseudo-VF morphogenesis. **(a)** Schematic representation of M2-1’s heterogeneous condensation with P on the first hand, and RNA on the second hand. Made with *Biorender.com* **(b)** Phase diagrams obtained in the presence of mCherry-P/WT-P and M2-1. **(c)** Phase diagrams obtained in the presence of mCherry-P/WT-P and N rings, with increasing concentrations of M2-1. The scatter plots display the approximate likelihood of phase separation. Red and blue points indicate phase separated and mixed regions respectively. The dashed line depicts the phase boundary (acting as visual reference). The number of distinct microdroplets examined **(b)** n= 140802 (left), n=61724 (right), **(c)** n= 31076 (left), n=31585 (centre), n=73793 (right). **(d)** Phase diagrams obtained in the presence of M2-1 and 5’ OH RNA-Cy5 (left) or 5’ cap RNA-Cy5 (right). Both are of the same sequence (GGGCAAAAGCGUAC), which consists of 7 nucleotides of the RSV gene start (GS) signal, followed by 7 random nt. **(e)** Analysis of M2-1 (left) or monomeric N-P40 (right) protein with 5’ OH or 5’-cap 7mer RNA (random sequence AGCGUAC) by agarose native gel band shift assay.

These results confirm the capacity of P and M2-1 to undergo phase separation and that M2-1 is recruited to N/P condensates. More interestingly, our data reveal that in contrast to the addition of monomeric N which inhibits LLPS, M2-1 actively facilitates condensate formation. These findings suggest that M2-1 participates in pseudo-VF morphogenesis by increasing the overall multivalency, thereby enhancing LLPS capacity.

### The 5’ cap of RNA favors M2-1 binding

Following its recruitment to VF, M2-1 binds to nascent mRNAs, leading to the formation of IBAG sub-compartments (*4*, *20*). While RNA sequence and length are known to be important factors for M2-1 binding, a definitive understanding of the elements driving this selective recognition of mRNAs over genomic RNA remains elusive (*19*, *30*, *32*, *40*, *41*). A major difference between (anti)genomic RNA and mRNA is the presence of a cap on the 5’ end of the latter, which consists of a methylated guanosine nucleotide (^7m^G) attached to the 5’ end of the RNA strand by a 5’-tri-phosphate bridge (ppp), as well as methylation of the 2’ oxygen (2’Ome) from the first 5’ nucleotide. To evaluate whether LLPS events between M2-1 and RNA might be influenced by such a difference in RNA nature, we generated phase diagrams for M2-1 in combination with Cy5-labelled 14-mer nucleotides (nt) RNA strands, either capped (5’ ^7m^GpppGm) or not (5’ OH) (**Figure 4d**). The length of these RNAs was chosen based on previous findings stating that 13-mer is the optimal length for M2-1 binding (*30*). Also based on previous RSV MTase assay (*42*), we used the RNA sequence GGGCAAAAGCGUAC-Cy5, consisting in 7 nt corresponding to the RSV gene start (GS) signal, followed by 7 random nt, with a Cy5 on the 3’ end. We observed that 5’ OH RNA readily phase separated with M2-1 across a wide range of concentrations, and that LLPS was unaffected by an excess of RNA (**Figure 4d**). In contrast, a more restricted phase separation was observed with 5’ capped RNA and M2-1, and LLPS was completely inhibited by an excess of 5’ capped RNA (**Figure 4d**). Given that LLPS is governed by weak, transient interactions, these results suggested that M2-1 interacts with a higher-affinity with capped RNA compared to uncapped RNA. To directly test the role of 5’ cap on M2-1 binding, we next analysed M2-1-RNA interactions by band shift on agarose native gel. Previous work showed that M2-1 possesses two RNA binding sites, is capable of binding RNA strands as short as 7-nt long, and has a higher affinity for specific sequences, such as the gene-end (GE) signal, than for random sequences (*19*, *30*, *32*). We therefore used a 7-nt random RNA sequence (ACGCGAA) to mimic a minimal binding event. This same RNA length was also found to be the minimum required for encapsidation by the N protein, used as a control (*34*). No band shift was observed for M2-1 upon incubation with the 5’ OH RNA, indicating that M2-1 did not bind this uncapped, short RNA strand (**Figure 4e**). However, a clear band shift was observed in the presence of the 5’ capped RNA, indicative of binding. In a stark contrast, the monomeric N-P40 protein was able to interact with the 5’ OH RNA but showed no binding to the 5’ capped RNA, in agreement with previous published data (**Figure 4e**) (*34*).

Our results demonstrate for the first time that the presence of a 5’ cap on RNA plays a critical role in the RNA-M2-1 interaction. Considering the strong RNA-binding affinity of both M2-1 and N, our observations offer a compelling model for their distinct roles within VF: the 5’ cap acts as a key determinant that directs M2-1 to nascent mRNAs while simultaneously creating a steric hindrance that prevents N from encapsidating them (**Figure 5**). This selectivity ensures that N is free to specifically encapsidate uncapped RNAs, i.e., the viral genome and antigenome.

**Figure 5.**
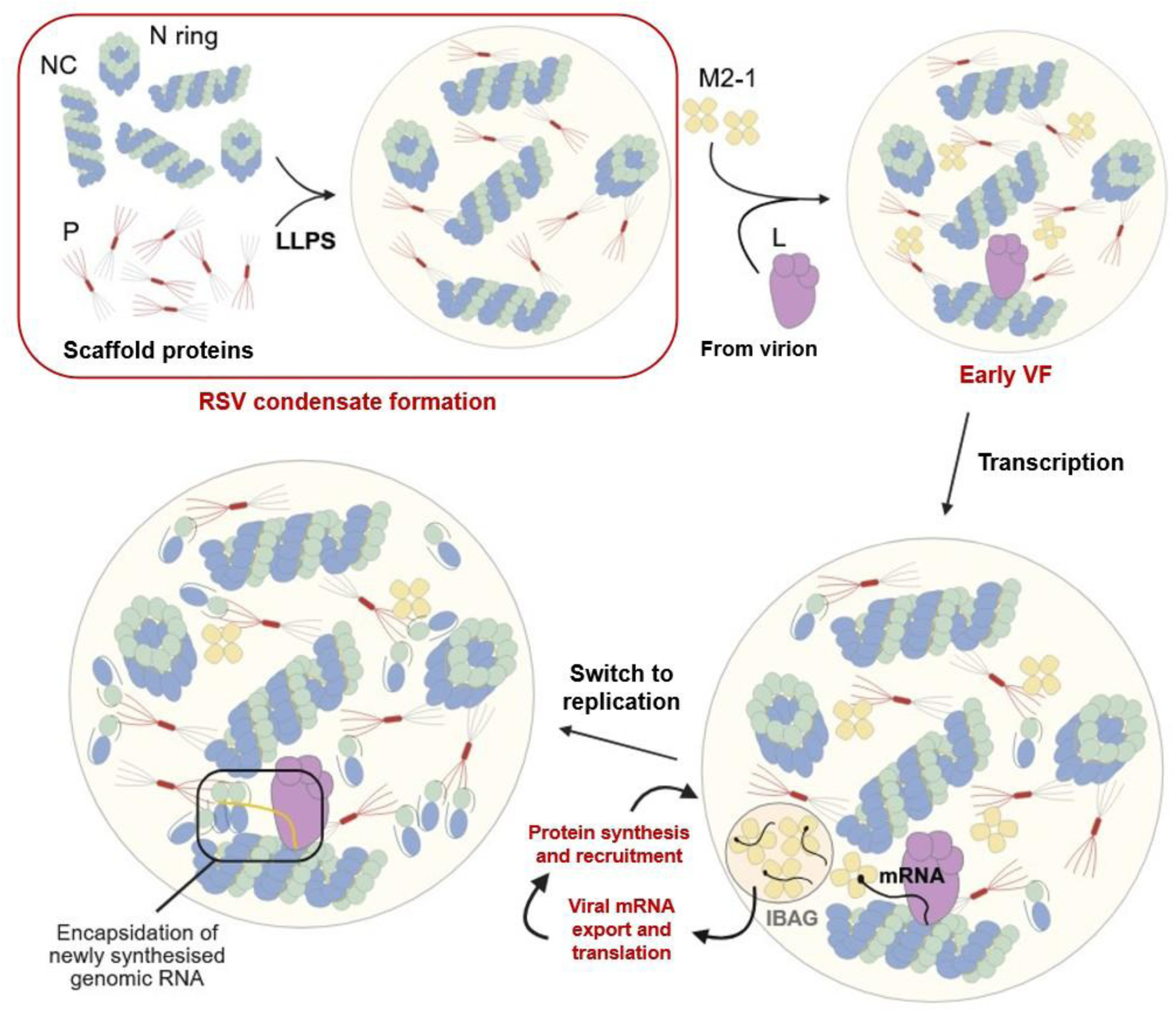
A comprehensive model of RSV VF morphogenesis and function. NC, nucleocapsid; LLPS, liquid-liquid phase separation; VF, viral factory; L, viral polymerase; mRNA, messenger RNA; IBAG, inclusion body associated granules. Made with *Biorender.com*.

## Discussion

As for most viruses from the *MNV* order, RSV replication and transcription take place within cytoplasmic VF formed upon LLPS induced by the viral N and P proteins. These membraneless organelles concentrate viral and cellular proteins, enabling efficient viral polymerase activity and regulation of host immune responses. Given their critical role in the viral life cycle and the absence of cellular counterparts, VF are attractive targets for antiviral development. For example, cyclopamine (CPM) has been shown to inhibit RSV replication by altering the interactions between M2-1 and P, which impairs protein mobility within VF and ultimately hardens the condensates (*38*, *43*). In this study, we set out to characterise the roles of the core viral proteins N, P and M2-1, as well as that of RNA, in the induction and regulation of LLPS events underlying VF formation and dynamics.

Using recombinant proteins without crowding agents, we showed that optimal N/P pseudo-VF formation occurs within a tightly controlled stoichiometric window of P tetramers per N ring. Analysis of our phase diagram showed that LLPS is bounded by a minimal P_tetramer_:N_ring_ threshold of approximately 0.5:1 to 1:1 and by a P-induced re-entry of phase at the high end, ranging from 3:1 to 12:1 (**Figure 2a**). This behaviour is strongly influenced by the absolute N concentration, emphasizing the role of the biochemical environment. This P-excess inhibition is corroborated by our cellular results: an excess of P impairs pseudo-VF morphogenesis and inhibits polymerase activity (**Figure 2e, f**). Differences in optimal N:P ratios for LLPS *in vitro* and *in cellula* were however observed, revealing the importance of the environment for N/P induced LLPS. Further investigation of the impact of increased concentrations of crowding agents on RSV LLPS should allow the identification of conditions that could better mimic the cytoplasmic environment. Our findings further confirm that RNA, rather than the sole oligomeric state of N, is essential for LLPS between N and P (**Figure 2b**). This aligns with structural data showing that RNA accessibility and the conformation of N protomers within nucleocapsid-like helices may impact LLPS (*18*). Although the study of recombinant nucleocapsids would have been highly relevant in the present study, their inherent fragility made them incompatible with the high-pressure and narrow channels of microfluidic chips used to perform phase scan experiments. We also explored how the concentration of specific proteins and their domains regulate LLPS. Our results with P domains confirm previous findings that the P C-terminal part and oligomerisation domain (P_127-241_) are sufficient to induce LLPS with N rings (**Figure 2c, d**) (*22*). Importantly, we found that P_127-241_ induced relatively higher levels of phase separation than full-length P without significantly affecting affinity to N, suggesting that the highly disordered P_NTD_ negatively affects LLPS, likely through steric hindrance caused by its unstable structure.

Our work sheds light on the dual role of the N protein pool. Using a three-component system, we showed that while N-P40 can be recruited into *in vitro* N/P condensates (*35*), an excess of it impairs LLPS (**Figure 3**). Surprisingly, native gel analyses also showed that although the interaction between P and N-P40 is weak, ternary N-P-N-P40 complexes can form. An excess of N-P40 increases the proportion of P-N-P40 and N-P-N-P40 complexes at the expense of N-P complexes, which can explain its negative impact on N/P LLPS (**Figure 3b**). Similarly, an artificial increase of the monomeric N-P40 pool in transfected cells down-modulated pseudo-VF morphogenesis, and inhibited viral polymerase activity (**Figure 3c, d**). Although complementary strategies using more relevant N^0^ surrogate (i.e., without P40 fusion) are needed to validate our observations, this suggests that regulation of the N^0^ pool may be important for VF formation and function. This would also be consistent with previous indications that the size of the N^0^ pool varies throughout the replication cycle and may induce a switch in L activities, from transcription to replication at later stages (*44*).

Finally, we explored the role of M2-1, which is recruited to early VF by P and later concentrates with nascent mRNAs within liquid-like subcompartments from which N, P, L and genomic RNA are excluded (*4*, *19*, *20*). We confirmed the capacity of P and M2-1 to phase separate without crowding agents and, importantly, we showed that an excess of M2-1 does not disrupt LLPS (**Figure 4b**). Unlike monomeric N, M2-1 actively facilitates LLPS with N rings and P tetramers (**Figure 4c**) (*37*, *38*), seemingly by increasing the system’s overall multivalency. The recruitment of M2-1 in VF involves a low nanomolar affinity interaction with the P_NTD_, theoretically too strong for phase separation (*39*). Interestingly however, M2-1 was recently shown to phase separate with the full length-P *in vitro*, but also with P deleted or either the P_NTD_ or P_CTD_, which suggests that other weaker interactions occur between several P domains and M2-1 tetramers (*37*). Our findings correlate with this projected multivalency. We propose that the addition of M2-1 increases the system’s multivalency and possibly stabilises P_NTD_, thereby up-regulating N/P LLPS. Overall, these results support a role of M2-1 in VF morphogenesis (*37*, *38*). Our most significant finding however, pertains to M2-1’s interaction with RNA, which may provide insights into the mechanisms underlying the presence of sub compartments where M2-1 and viral mRNA concentrate within VF during RSV infection. We observed for the first time that the nature of the RNA influenced LLPS with M2-1. More specifically, we showed that M2-1 preferentially binds to 5’ capped RNA over uncapped RNA (**Figure 4d, e**), contrary to N^0^ which encapsidates short, uncapped RNA but not 5’ capped ones (**Figure 4e**) (*34*). Our data provide a compelling model for the presence of subcompartments within VF and for the specific encapsidation of genomes and antigenomes by N, governed respectively by the presence or lack of a 5’ cap (**Figure 5**). This new data reinforces the idea that the specific recognition of nascent mRNA by M2-1 within VF may influence the encapsidation of genomes and antigenomes by N^0^. Resolving the crystal structure of an M2-1 tetramer bound to capped RNA will be essential to precisely define the molecular basis for this specific recognition.

In conclusion, this study highlights new information on the roles of N, P, M2-1 and RNA on RSV pseudo-VF morphogenesis and dynamics (**Figure 5**). Although several other phase scan combinations could have provided more information, for instance N rings with P, M2-1 and capped RNA to define optimal ranges for the formation of pseudo-VF sub compartments, such complexification of the system is not feasible yet. Involving additional components in the future, such as the viral polymerase L, or cellular factors, should also provide critical insights into LLPS. This work also highlights the potential of exploring LLPS in other viral families, to reveal key insights into these fascinating processes, and to identify novel antiviral targets.

## Author contributions

V.B., T.A., J.-F.E. and M.G. designed experiments. V.B., C.-A.R. and M.G. performed proteins’ purification. T.A. and V.B. performed the *in vitro* phase separation assays. J.T., J.-J. V. and F. D. synthesised RNAs. V.B. and M.G. performed the cellular experiments. E.C. performed the MDS assays. V.B. wrote the paper with contributions from all authors, and M.G. edited the manuscript. All authors commented on the manuscript.

## EXPERIMENTAL PROCEDURES

### Cells

BHK-21 cells (clone BSRT7/5) constitutively expressing the T7 RNA polymerase (*45*) and BEAS-2B cells (ATCC CRL-9609) were grown in Dulbecco’s modified Eagle’s medium (Eurobio Scientific, Les Ulis, France) supplemented with 10% fetal calf serum (FCS) (Eurobio), 1 mM L-Glutamine, and antibiotics. The transformed human bronchial epithelial cells BEAS-2B (ATCC CRL-9609) were maintained in Roswell Park Memorial Institute (RPMI) 1640 medium (Eurobio) supplemented with 10% FCS, 1% L-glutamine, and antibiotics. The cells were grown at 37°C in 5% CO_2_ and were transfected using Lipofectamine 2000 (Invitrogen) as described by the manufacturer.

### Minigenome assay

BSRT7/5 cells at 90% confluence in 96-well dishes were transfected with a plasmid mixture containing 62.5 ng of pM/Luc, 62.5 ng of pN, 62.5 ng of pP (or different ratios of pP, pN, and pN-P40 plasmids), 31.25 ng of pL, and 15.5 ng of pM2-1 as well as 15.5 ng of pRSV β-gal (Promega) to normalise transfection efficiencies (*46*). For each experiment, 5 wells were transfected and independent experiment were performed three times. Cells were lysed 24 hours after transfection in luciferase lysis buffer (30 mM Tris, pH 7.9, 10 mM MgCl2, 1 mM DTT, 1% Triton X-100, and 15% glycerol). The luciferase activities were determined for each cell lysate with an Infinite 200 Pro (Tecan, Männedorf, Switzerland) and normalised based on β-galactosidase (β-gal) expression.

### Plasmid constructs

For expression and purification of N recombinant proteins, the previously described pET-N, pET-mCherry-N, pGEX-PCT, and pET-N-P40 plasmids were used (*18*, *23*, *34*). For expression and purification of P recombinant proteins, the previously described pGEX-P, pGEX-mCherry-P, pGEX-P[1-126], and pGEX-P[127-241] plasmids were used (*22*). For expression and purification of M2-1 recombinant proteins, the previously described pGEX-M2-1 plasmid was used (*19*, *20*, *38*).

pcDNA3.1 codon-optimized plasmids for mammalian expression encoding the RSV A2 P, N, and N-P40 proteins were already described (*35*, *47*). The plasmids used for the minigenome assay were also previously described (*46*).

### Expression and purification of the recombinant proteins

The *Escherichia coli* BL21 (DE3) bacteria strain (Novagen, Madison, WI) were transformed with the different mentioned plasmids. Production and purification of the recombinant proteins was conducted as previously described (*19*, *20*, *22*, *34*, *38*).

### Synthesis of RNA substrates

RNA substrates with the sequence (GGGCAAAAGCGUAC) were chemically synthesised on LCAA-CPG solid support (Biosearch Technologies) at 1 µmole scale using an ABI 394 automated synthesiser (Applied Biosystems) with TWIST synthesis columns (Glen Research) and oligonucleotide synthesis reagents from Biosearch Technologies. Building blocks 2’-*O*-pivaloyloxymethyl (PivOM) or 2’-*O*-propionyloxymethyl (PrOM) 3’-*O*-phosphoramidite ribonucleosides and 2’-*O*-methyl 3’-*O*-phosphoramidite guanosine to obtain ^7m^GpppG_m_-RNA were purchased from ChemGenes Corp., USA. 3’-cyanine5 CPG was obtained from Biosearch Technologies. After RNA assembly, depending on the desired 5’ end of the RNA, different treatments were applied. For 5’ OH-RNA sequence (GGGCAAAAGCGUAC-Cy5) prepared from 2’-*O*-PivOM nucleotides (*48*), the solid support was treated with a solution of 1M DBU/CH_3_CN (2 mL) for 3 min at room temperature. Then, RNA was cleaved from solid support using a 28% aqueous ammonia solution at 40°C for 3 h. The solvents were evaporated under vacuum (in the presence of 500 µL of *iso*propylamine). For 5’ ^7m^GpppG_m_GGCAAAAGCGUAC-Cy5, prepared from 2’-*O*-PrOM nucleotides and 2’*O*Me G (*49*), after elongation the 5’-hydroxyl group was phosphorylated and the resulting *H*-phosphonate derivative was oxidised and activated to a phosphoroimidazolidate derivative to react with ^7m^GDP, yielding ^7m^Gppp-G_m_ RNA, Solid support was treated with a solution of 1M DBU/CH_3_CN (2 mL) for 3 min at r.t and RNA was released from CPG and deprotected using a 7M methanolic ammonia solution for 3 h at 40 °C. Solvents were evaporated under vacuum (in the presence of 500 µL of *iso*propylamine). RNA substrates were analysed and purified by IEX-HPLC (*>* 95% pure) and were characterised by MALDI-TOF mass spectrometry. RNA substrates were desalted using a C_18_ cartridge Sep-Pak® Classic, lyophilised and stored at -20 °C.

### Fabrication of microfluidic devices

Microfluidic device designs were created using AutoCAD software and fabricated via standard soft-photolithography techniques. The fabrication process employed SU-8 photoresist patterned on silicon wafers to produce PDMS-on-glass microfluidic chips, as previously described (*50*–*52*). Briefly, SU-8 3050 (A-Gas Electronic Materials Limited) was poured onto polished silicon wafers (MicroChemicals GmbH) and spin-coated at 3000 RPM for 45 seconds. The coated wafers were then subjected to a soft bake on a level hot plate at 95°C for 15 minutes. Following this, an acetate mask sheet containing the microfluidic layout was aligned and placed onto the SU-8-coated wafer and exposed to UV light for 40 seconds at room temperature. The mask was removed immediately post-exposure, and the wafer underwent a post-exposure bake at 95°C for 5 minutes. Development was carried out in a PGMEA (propylene glycol monomethyl ether acetate; Sigma-Aldrich) bath for 10–15 minutes with gentle agitation to eliminate unexposed photoresist. The wafers were then rinsed with isopropyl alcohol and dried using a nitrogen stream. This process yielded master molds with microchannel features approximately 50 μm in height. Poly (dimethyl siloxane) (PDMS; Sylgard 184 kit; Dow Corning) and crosslinking agent was thoroughly mixed in a ratio of 10:1. The mixture was poured over the master mold placed in a plastic petri dish and cured at 60 °C for 2 hours. Once cured, individual PDMS devices were cut using a scalpel, and inlet and outlet holes were punched. The cut PDMS slabs were then cleaned by immersion in isopropyl alcohol (IPA) and subjected to sonication for 15 minutes and dried with nitrogen stream. To assemble the microfluidic device, the PDMS layer was bonded to a pre-cleaned 1 mm thick glass slide (Epredia) using oxygen plasma treatment (Diener Femto Electronics, 40% power, 30 seconds). For hydrophobic surface modification, the microchannels were treated with a 1% v/v solution of trichloro(1H,1H,2H,2H-perfluorooctyl) silane (Sigma-Aldrich) in HFE-7500 fluorinated oil (3M™ Novec™ Engineered Fluid) for 2 minutes. This was followed by immediate drying on a flat hot plate at 95 °C for 10 minutes. Finally, the channels were flushed with HFE-7500 and dried with a nitrogen stream.

### Generation of phase diagrams

Phase scan, a semi-automated droplet-based combinatorial microfluidic platform, was employed to generate and analyse multidimensional 2D and 3D phase diagrams, as described previously (*33*). In a typical Phase scan run, three-component (e.g., N, P, and buffer) and four-component (e.g., N, P, M21, and buffer) aqueous mixtures were used for 2D and 3D diagrams, respectively. A microfluidic flow controller (Flow EZ™, flow unit, OxyGEN; Fluigent) facilitated pressure-based control of both aqueous and oil inputs. Fluorescently labeled aqueous components—such as N (Alexa Fluor 488 carboxylic acid), P (mCherry), and, when applicable, M21 (Alexa Fluor 647 carboxylic acid; ThermoFisher Scientific)—were used alongside unlabelled buffer, with each component (except buffer) uniquely barcoded using a distinct fluorophore depending on the experiment. A pre-programmed flow profile automatically modulated the input flow rates of aqueous solutions to achieve precise combinatorial mixing. For 2D PhaseScan cycles, the total aqueous flow rate was set to 60 µL/h, with individual component flow rates ranging from 5 to 50 µL/h. For 3D cycles, a total aqueous flow rate of 80 µL/h was used, with individual flow rates ranging between 5 and 65 µL/h. The oil phase HFE-7500 fluorinated oil supplemented with 1.2% w/v fluorosurfactant (RAN Biotechnologies) was introduced at a constant flow rate ranging from 50-100 µL/h to enable droplet formation in the microfluidic device. Droplets were imaged under continuous flow using an epifluorescence microscope (Cairn Research) equipped with a 10x objective lens (Nikon CFI Plan Fluor 10x, NA 0.3). Image analysis was carried out using a custom Python script. Droplets were identified and fitted as square regions in the acquired images, and any false detections were excluded. Total fluorescence intensity within each fitted area was calculated, normalised for droplet volume (based on estimated diameter), and converted into concentrations using calibration data from reference samples. Droplets were then classified as either phase-separated or mixed based on the presence or absence of visible condensates. The data was represented graphically as a scatter plot with an overlaid color-coded indicating the approximate likelihood of phase separation.

### Band shift on native agarose gels

Samples of P, N, N-P40 proteins alone or mixed, or of RNA oligonucleotides (15 µM) and purified N-P40 or M2-1 proteins (10 µM) were incubated for 30 min at room temperature in 20 mM Tris pH8 150 mM NaCl and further analysed by band shift on agarose native gel. 50% sucrose loading buffer was added to the samples before loading on native 1% agarose gel. Migration was performed in 1X Tris-Glycine buffer during 1 h 30 at 80 V before staining with amido black 10B.

### Immunofluorescence

BEAS-2B cells grown on coverslips in p24 well-plates were transfected with different ratio of pcDNA-N and pcDNA-P (0.8 µg of DNA total per well), or with pcDNA-P (0.4 µg) and different ratio of pcDNA-N and pcDNA-N-P40 (up to 04. µg of DNA). Twenty-four hours after transfection, cells were fixed with 4% paraformaldehyde (PFA) for 20 min. Fixed cells were permeabilized, blocked for 30 min with phosphate-buffered saline (PBS) containing 0.1% Triton X-100 and 3% bovine serum albumin (BSA), and then successively incubated for 1h at room temperature with primary (rabbit anti-N antiserum (*53*) and mouse anti-P (*54*), or rabbit anti-P antiserum (*46*), and secondary antibody (mouse and rabbit IgG coupled to Alexa 568 or Alexa 488 (Invitrogen) mixtures diluted in PBS containing 3% BSA. For nuclei labelling, Hoechst 33342 (Invitrogen) was added during incubation with the secondary antibodies. Coverslips were mounted in Prolong gold antifade reagent (Invitrogen). Cells were observed with a Nikon TE200 microscope equipped with a CoolSNAP ES² (Photometrics) camera, and images were processed using MetaVue (Molecular Devices) and ImageJ software.

### MDS

Microfluidic diffusional sizing of the samples was carried out as previously described (*55*). In brief, the channels of a Fluidity One-M microfluidic plate (Fluidic Sciences) were prefilled with buffer (150 mM NaCl, 20 mM TRIS, pH7.4). Following this, samples containing 0.5 µM N-mCherry and increasing concentrations of unlabelled P protein or P domains (incubated in the dark, 30 mins) were loaded into the sample channels. The average hydrodynamic radius of N was measured in triplicate using Alexa-647 detection setting and size-range setting of 4.7–20 nm was used and binding affinities extracted from the fit.

### Funding information

This work was carried out with the financial support of the French Agence Nationale de la Recherche, specific program ANR RSVFact (ANR-21-CE15-0030-02), ANR SofteN (ANR-23-CE11-0027-01) and ANR DecRisP (ANR-19-CE11-0017-01). We would like to acknowledge funding from the European Research Council under the European Union’s Seventh Horizon 2020 research and innovation program through the ERC grant DiProPhys [101001615] (T.A. and TPJK), and ERC ECH2020 through the Marie Skłodowska-Curie grant MicroREvolution (agreement no. 101023060 to T.S.), the Frances and Augustus Newman Foundation (T.S., EDC), and the Finlay family (T.S.).

### Conflict of interest

The authors declare that they have no competing interests.

### Data availability

All data needed to evaluate the conclusions in the paper are present in the paper and/or the Supplementary Materials.

## Notes

### Competing Interest Statement

The authors have declared no competing interest.

