## Supplemental Figures for "*In vitro* liquid-liquid phase separation induced by respiratory syncytial virus proteins and RNA"

Supplementary Information

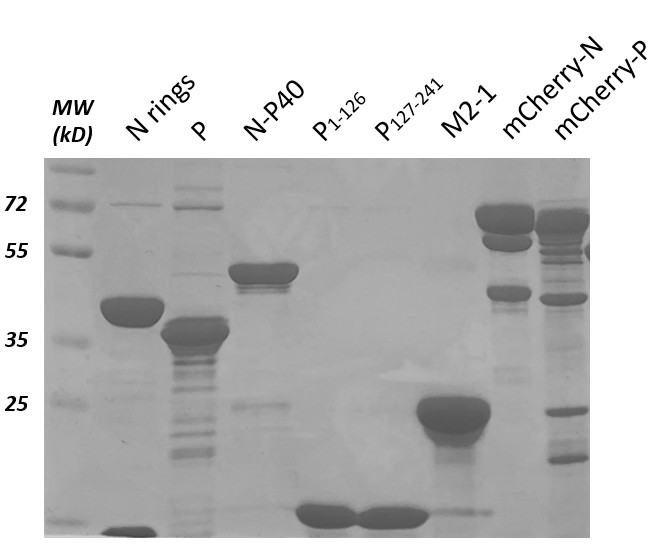
**Figure S1 – SDS-PAGE of the recombinant proteins used in the present study.**

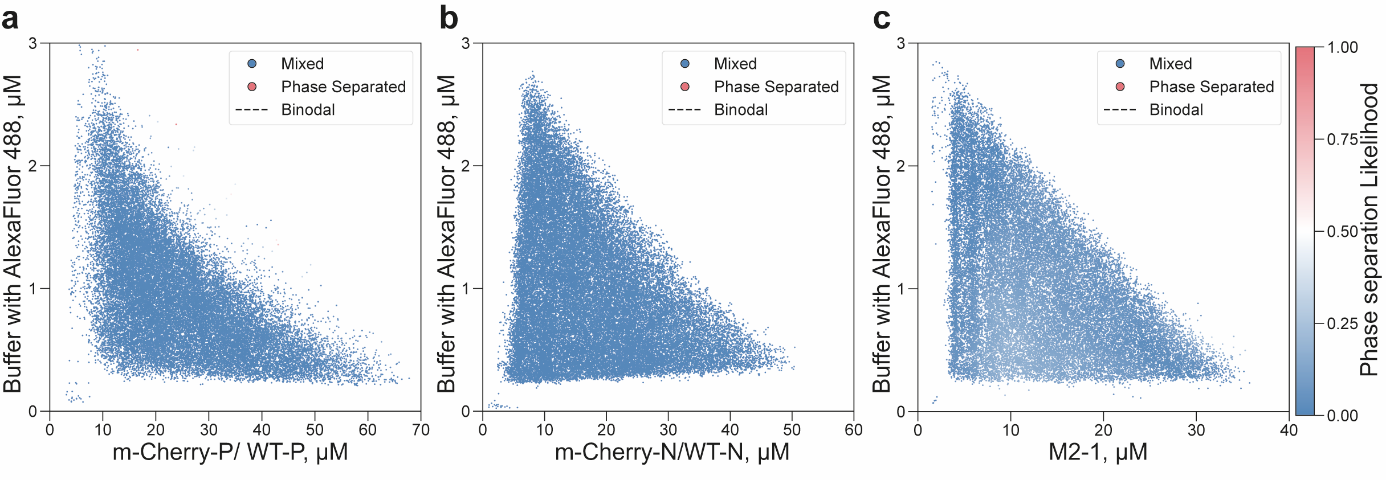

**Figure S2 – Control phase scan experiments.** Phase diagrams obtained in the presence of **(a)** mCherry-P/WT-P, n = 34534, **(b)** mCherry-N/WT-N rings, n = 44975, and **(c)** M2-1, n = 34669, alone at different concentrations. The scatter plot displays the approximate likelihood of phase separation. Red and blue points indicate phase separated and mixed regions respectively. No phase separation was observed.

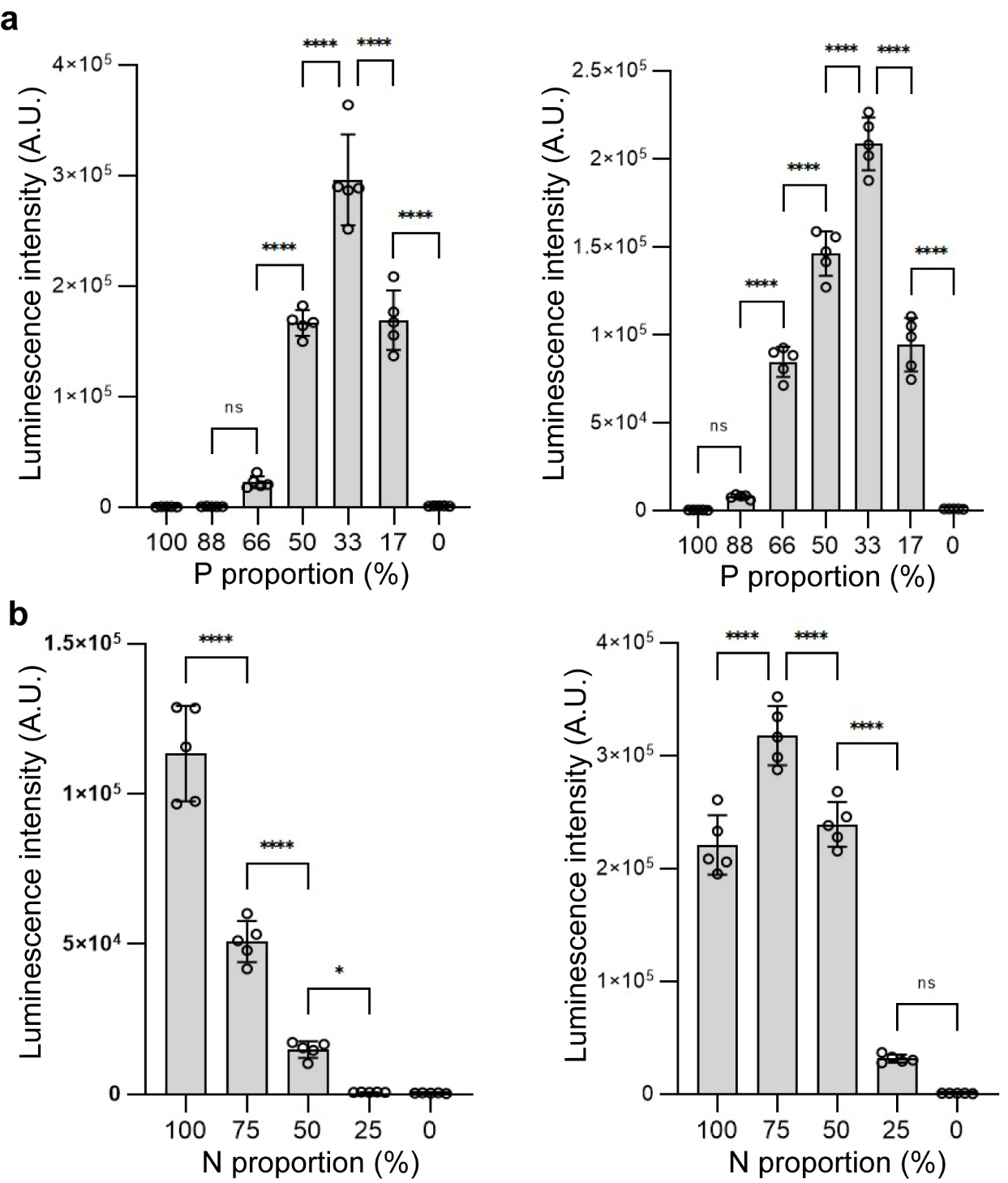

**Figure S3 – Replicate minigenome activity assays. (a)** BSRT7/5 cells were transfected with plasmids encoding viral proteins M2-1 and L, varying quantities of plasmid DNA encoding N and P, a plasmid encoding the pMT/Luc minigenome as well as the pCMV-βGal for transfection standardisation. Cells were lysed 24h post-transfection and viral RNA synthesis was quantified by measuring the luciferase activity. Each luciferase minigenome activity value was normalised based on β-galactosidase expression. For each condition, five wells were transfected. **(b)** BSRT7/5 cells were transfected with plasmids encoding P, M2-1, L, pMT/Luc minigenome, pCMV-βGal, and varying quantities of N and N-P40. Cells were lysed 24h post-transfection and viral RNA synthesis was quantified by measuring the luciferase activity. Each luciferase minigenome activity value was normalised based on β-galactosidase expression. For each condition, five wells were transfected.
